# Parallel evolution of sperm hyper-activation Ca^2+^ channels

**DOI:** 10.1101/120758

**Authors:** Jacob C. Cooper, Nitin Phadnis

## Abstract

Sperm hyper-activation is a dramatic change in sperm behavior where mature sperm burst into a final sprint in the race to the egg. The mechanism of sperm hyper-activation in many metazoans, including humans, consists of a jolt of Ca^2+^ into the sperm flagellum via CatSper ion channels. Surprisingly, *CatSper* genes have been independently lost in several animal lineages. In *Drosophila*, sperm hyper-activation is performed through the co-option of the polycystic kidney disease 2 (*Dpkd2*) Ca^2+^ channel. The parallels between *CatSpers* in primates and *Dpkd2* in *Drosophila* provide a unique opportunity to examine the molecular evolution of the sperm hyper-activation machinery in two independent, nonhomologous calcium channels separated by more than 500 million years of divergence. Here, we use a comprehensive phylogenomic approach to investigate the selective pressures on these sperm hyper-activation channels. First, we find that the entire *CatSper* complex evolves rapidly under recurrent positive selection in primates. Second, we find that *pkd2* has parallel patterns of adaptive evolution in *Drosophila*. Third, we show that this adaptive evolution of *pkd2* is driven by its role in sperm hyper-activation. These patterns of selection suggest that the evolution of the sperm hyper-activation machinery is driven by sexual conflict with antagonistic ligands that modulate channel activity. Together, our results add sperm hyper-activation channels to the class of fast evolving reproductive proteins and provide insights into the mechanisms used by the sexes to manipulate sperm behavior.

## Introduction

Sexual conflict shapes sperm development and sperm dynamics (Swanson and Vacquier 2002; Clark et al. 2006; Turner et al. 2008; Wilburn and Swanson 2016). Both male-female interactions and inter-male competition drive rapid changes in male reproductive proteins, whose constant innovation has been likened to a molecular arms race. These rapid changes in reproductive proteins have the potential to establish barriers to fertilization between populations and lead to the evolution of new species (Parker and Partridge 1998; Gavrilets 2000; Howard et al. 2009; Moyle et al. 2014). The best-known examples of this phenomenon include the rapid evolution of reproductive proteins in abalone, mammals, and *Drosophila* (Lee et al. 1995; Swanson and Vacquier 1997; Kresge et al. 2001; Swanson et al. 2003; Clark et al. 2007; Hamm et al. 2007; Findlay et al. 2014; Vicens et al. 2015). Molecular evolutionary studies on how sexual conflicts shape reproductive proteins have focused on several aspects of sperm biology such as direct sperm-egg interactions, seminal fluid proteins and sperm behavior (Swanson and Vacquier 2002; Panhuis et al. 2006; Fisher et al. 2016). Here, we uncover the patterns of molecular evolution of the sperm hyper-activation machinery in animals, which remains a fundamental but largely unexplored aspect of sperm biology.

When spermatogenesis is complete, the resulting mature sperm are motile but quiescent. After copulation, however, sperm cease normal swimming and burst into a sprint. This dramatic post-mating acceleration of sperm is known as sperm hyper-activation (Suarez et al. 1991). Sperm hyper-activation was first observed in mammals through studies on the golden hamster (Yanagimachi 1969; Yanagimachi 1970), and has since been since described in many other taxa (Suarez and Ho 2003; Cosson et al. 2008). When sperm hyper-activate, they go through a cellular change that alters the motion of the sperm flagellum from a slow, low amplitude, symmetric beat to a whip-like, high amplitude, asymmetric beat (Ooi et al. 2014). This transition is not subtle; hyper-activated sperm swimming at top speed are propelled with a force several times that of normal swimming (Ishijima 2011). This hyper-activated acceleration of sperm is necessary for successful fertilization, and is an integral part of sperm capacitation.

The proximate molecular mechanism of sperm hyper-activation consists of a jolt of Ca^2+^ ions to the sperm flagellum, which triggers a complex intracellular chain of events to drive accelerated swimming (Ho et al. 2002; Carlson et al. 2009). The use of Ca^2+^ influx as a sperm hyper-activation trigger is an ancient and widely conserved mechanism across metazoans (Cai et al. 2014). In most metazoans, the Ca^2+^ influx arrives via the <u>ca</u>tion channels of <u>sper</u>m (CatSper) complex, which forms ion channels on the sperm flagellum (Cai et al. 2014). The CatSper complex consists of four core proteins that form the Ca^2+^ pore and five auxiliary proteins (Chung et al. 2017). Sperm that do not hyper-activate fail to fertilize eggs, and males with mutations that disable sperm hyper-activation are often sterile despite producing morphologically normal sperm (Ren et al. 2001).

All CatSper proteins–core and auxiliary–are necessary for sperm hyper-activation and male fertility (Loux et al. 2013). Despite the conserved role of the CatSper complex in sperm hyper-activation across metazoan taxa, the sequences of CatSper proteins provide hints of being evolutionarily labile. Previous analyses of *CatSper* evolution have been focused on the first exon of *CATSPER1*, which shows signs of adaptive evolution in the form of an accelerated accumulation of indels and non-synonymous changes in mammals (Podlaha and Zhang 2003; Podlaha et al. 2005; Cai and Clapham 2008; Vicens et al. 2014). More surprisingly, the entire suite of *CatSper* genes has been lost in several animal taxa, including those leading to arthropods, nematodes, mollusks, jawless fishes, bony fishes, birds, frogs, etc. (Cai and Clapham 2008). Some taxa that have lost the *CatSper* complex, such as frogs, no longer hyper-activate their sperm during fertilization (Dziminski et al. 2009). In contrast, many other taxa, such as flies and birds, are known to hyper-activate sperm despite lacking a functional *CatSper* complex (Köttgen et al. 2011; O’Brien et al. 2011; Yang and Lu 2011; Nguyen et al. 2014; Zhou et al. 2015). These patterns suggest that some taxa that are missing the CatSper complex may have compensated for the loss through the co-option of other mechanisms to perform sperm hyper-activation. The evolutionary forces that drive the repeated turnover of the sperm hyper-activation machinery remain unaddressed.

The mechanism of CatSper-independent sperm hyper-activation remains unknown in many taxa, but is best understood in *Drosophila melanogaster*. None of the *CatSper* genes are present in the *D. melanogaster* genome. Yet, *Drosophila* males hyper-activate their sperm post-copulation as a necessary step for successful fertilization. CatSper-independent sperm hyper-activation in *Drosophila* is performed by the protein polycystic kidney disease 2 (*Dpkd2*). Similar to the CatSper proteins in mammals, *Dpkd2* is a Ca^2+^ ion channel protein on the fly sperm flagellum. *Dpkd2* null sperm are morphologically normal, but do not hyper-activate after transfer to the female reproductive tract. As a result, *Dpkd2*-deficient sperm fail to reach the storage organs and are not retained in the female (Gao et al. 2003; Watnick et al. 2003). *Drosophila*, therefore, appear to have compensated for the loss of CatSper ion channels by using the Dpkd2 channel to trigger sperm hyper-activation.

Sperm hyper-activation is a powerful and tightly controlled behavioral switch for the final sprint in the race to the egg (Montoto et al. 2011). The sperm hyper-activation machinery may, therefore, be vulnerable to the pressures of both female choice and inter-male competition. An evolutionary arms race over the modulation of sperm hyper-activation can manifest as the rapid evolution of sperm hyper-activation channels. The parallels between *CatSpers* in primates and *Dpkd2* in *Drosophila* provide a unique opportunity to examine the how selection has shaped sperm hyper-activation machinery in two independent, non-homologous calcium channels. Here, we use a comprehensive phylogenomic approach with *CatSper* in primates and *pkd2* in *Drosophila* to investigate the selective pressures on the Ca^2+^ channels required for sperm hyper-activation. First, we find that all core and auxiliary proteins of the *CatSper* complex evolve rapidly under recurrent positive selection in primates. Second, we find that *pkd2* has similar patterns of positive selection in *Drosophila*, including increased amino-acid substitution and an accumulation of indels. Third, we show that the selective pressures of *Drosophila pkd2* and primate *PKD2* are radically different; primate *PKD2* is highly conserved and is not involved in sexual conflict. This provides a unique example where an otherwise slow-evolving ‘housekeeping’ gene is dragged into an evolutionary conflict and experiences adaptive evolution. Together, our study provides the first comprehensive analysis of the molecular evolutionary patterns of the sperm hyper-activation Ca^2+^ channels in primates and flies.

## Results

### The entire CatSper complex evolves adaptively in primates

Despite the critical role of the *CatSper* complex in sperm hyper-activation across a wide variety of metazoa, little is known about its molecular evolution. At the core of the CatSper complex lies a Ca^2+^ pore composed of a hetero-tetramer of CATSPER1-4. Each core CatSper protein contains a six-pass transmembrane domain with polycystic kidney disease (PKD) domains (Quill et al. 2001; Ren et al. 2001; Lobley et al. 2003; Jin et al. 2005; Qi et al. 2007: 20) (Figure 1B). In contrast, four of the five auxiliary proteins, CATSPERβ, CATSPERΔ, CATSPERγ, and CATSPERε, have a large extracellular region with one or two transmembrane domains (Liu et al. 2007; Wang et al. 2009; Chung et al. 2011; Chung et al. 2017). The fifth auxiliary protein, CATSPERζ, is a small intracellular scaffold that helps assemble the complex (Chung et al. 2017). The CatSper channel is, in theory, well positioned to be acted upon by sexual selection. It is found only on the sperm flagellum, with a substantial portion exposed to the external environment of the sperm, and its only known function is in fertilization. We were, therefore, interested in a comprehensive understanding of the molecular evolutionary patterns of the *CatSper* complex and in uncovering the evolutionary forces that drive the changes in these genes.

**FIG 1.**
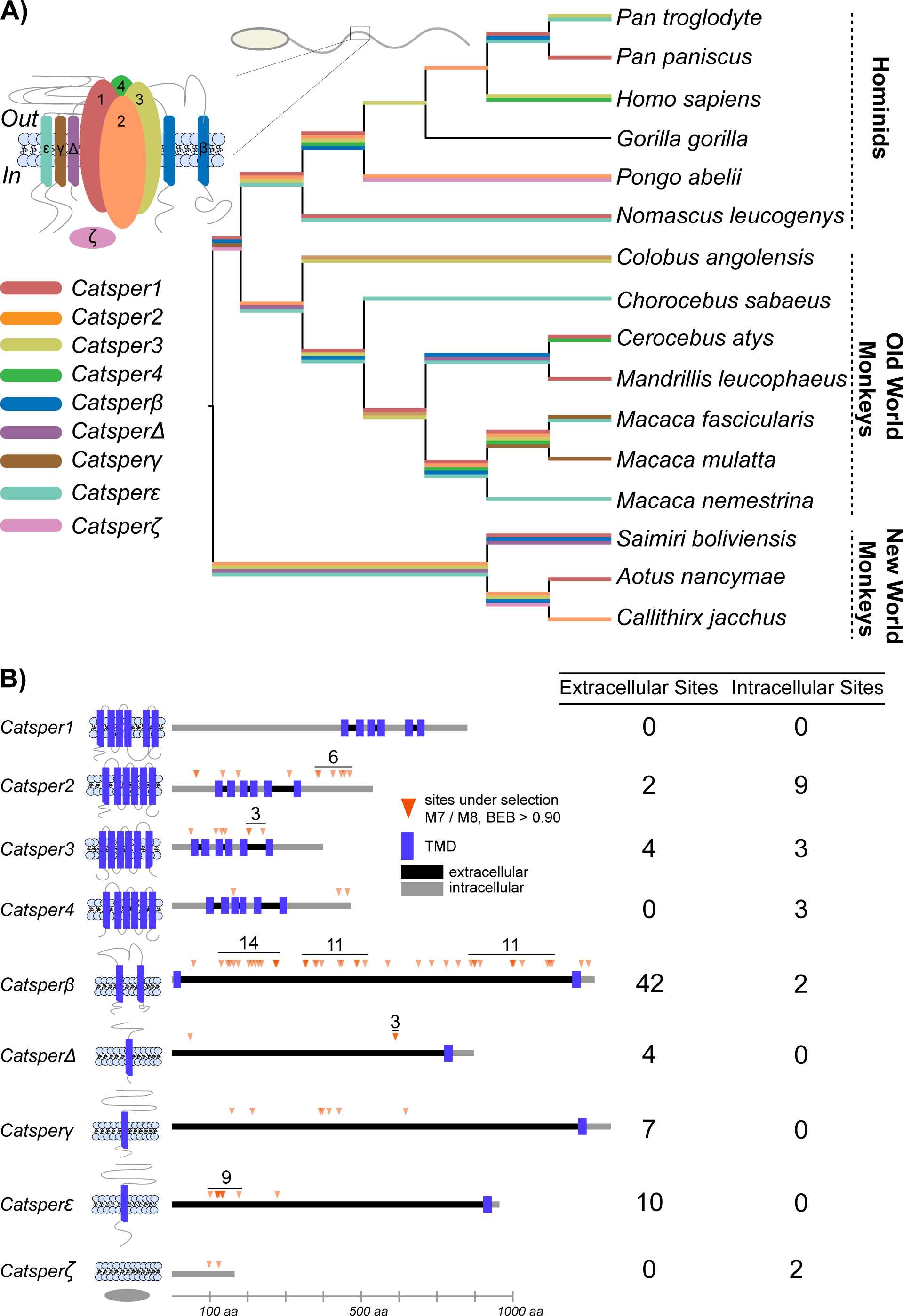
The entire *CatSper* complex evolves adaptively in primates. A) The *CatSper* complex evolves under positive selection in almost every lineage in the primate phylogeny. For each *CatSper* gene, branches where *dN/dS* >1 are highlighted with a color corresponding to each gene. In cases where multiple genes have a *dN/dS* >1 for a single branch, the colors are stacked. B) We detect specific sites under selection in almost every *CatSper* gene. Extracellular domains are marked as dark segments and intracellular domains as light segments. Orange arrows indicate the sites under selection by the Bayes-Empirical-Bayes test in PAML, with a posterior-probability greater than 0.90 (Yang et al. 2005). Selected sites grouped close together are labeled with a bar specifying the number of sites under selection. The number of extracellular and intracellular sites under selection for each gene are tabulated.

To examine the evolutionary forces that shape the CatSper complex in primates, we gathered and aligned homologous sequences for each *CatSper* gene from 16 primate species (Table 1). This sampling of primate species represents ~40 million years of divergence. To test if any of any of the CatSper proteins show signatures of positive selection, we used Phylogenetic Analysis by Maximum Likelihood (PAML) to calculate the synonymous to non-synonymous substitution rate ratios (*dN/dS*) (Yang 2007). For each of the nine genes that constitute the *CatSper* complex, we asked if any branch in the primate phylogeny had a *dN/dS* greater than 1. We first analyzed the four core proteins CATSPER1-4 that form the Ca^2+^ pore. We detected elevated rates of *dN/dS* in each of these four core proteins (Figure 1A; Supplementary Figure 1). In addition to the core proteins, all five auxiliary transmembrane proteins CATSPERβ, CATSPERΔ, CATSPERγ, CATSPERε and CATSPERζ also show strong signatures of adaptive evolution. We found that every branch in the primate lineage, with the exception of gorillas, shows a signature of positive selection for at least one of the *CatSper* genes. These results show that all nine proteins that form the CatSper complex have evolved under pervasive and strong positive selection across 40 million years of primate evolution.

**Table 1.**

| A) Primates | Group | B) Drosophila | subgroup |
| --- | --- | --- | --- |
| *Pan troglodyte* | Hominid | *D. melanogaster* | melanogaster |
| *Pan paniscus* | Hominid | *D. simulans* | melanogaster |
| *Homo sapiens* | Hominid | *D. sechellia* | melanogaster |
| *Gorilla gorilla* | Hominid | *D. mauritiana* | melanogaster |
| *Pongo abeli* | Hominid | *D. yakuba* | melanogaster |
| *Nomascus leucogenys* | Hominid | *D. tessieri* | melanogaster |
| *Colobus angolensis* | Old World Monkeys | *D. santomea* | melanogaster |
| *Chorocebus sabaeus* | Old World Monkeys | *D. erecta* | melanogaster |
| *Cerocebus atys* | Old World Monkeys | *D. orena* | melanogaster |
| *Mandrillus leucophaeus* | Old World Monkeys | *D. eugracillis* | eugracillis |
| *Macaca fascicularis* | Old World Monkeys | *D. biarmipes* | suzukii |
| *Macaca mulatta* | Old World Monkeys | *D. takahashii* | takahashii |
| *Macaca nemestrina* | Old World Monkeys | *D. ficusphila* | ficusphila |
| *Saimiri boliviensis* | New World Monkeys | *D. rhopaloa* | montium |
| *Aotus nancymae* | New World Monkeys | *D. elegans* | elegans |
| *Callithrix jacchus* | New World Monkeys | *D. ananassae* | ananassae |
|  |  | *D. bipectinata* | ananassae |

### Patterns of positively selected sites in CatSper channel proteins

The positions of the adaptively evolving amino acid sites within a protein can provide insights into the functional properties that are under selection. To identify the adaptively evolving amino acid sites in each of the *CatSper* genes, we used the NSsites models of PAML. For each *CatSper* gene, we tested whether its evolution over the primate phylogeny is consistent with neutral evolution (model M7 and M8a) or with recurrent positive selection (model M8). PAML also uses a Bayesian framework to identify the specific sites in a gene that evolve adaptively under recurrent positive selection. We found that all nine *CatSper* genes show significant evidence of recurrent positive selection with both of these tests (Supplementary Table 1). In CATSPER1, the site models of PAML did not detect any specific sites under selection, indicating that the selective signature may be broadly dispersed across the gene. *CATSPER2, CATSPER3*, and *CATSPER4* each have several sites that evolve under positive selection, but these are not clustered in any particular functional domain (Figure 1B; Supplementary Table 2). The patterns of selection in the auxiliary genes, however, are remarkably different. *CATSPERΔ, CATSPERγ*, and *CATSPERε* all have several sites under selection, and *CATSPERβ* has a dramatic excess of sites under selection compared to any other *CatSper* gene (Supplementary Table 2). The extracellular domain CATSPERβ is full of adaptively changing amino acid sites. Because *CATSPERβ* is such a clear outlier with the greatest number of sites under positive selection, we were concerned about spurious false positives generated from alignment errors. A manual inspection of the *CATSPERβ* alignment makes it clear that these are not false positives – we find practically no mis-alignment between the 16 homologous protein sequences (Supplementary Figure 2). Little is known about the precise molecular properties of CATSPERβ, other than that it requires CATSPER1 for stable localization to the tail of sperm (Liu et al. 2007). These results show that among all components of the CatSper complex, CATSPERβ is the most frequent target of adaptive evolution.

If the evolution of the sperm hyper-activation machinery is driven by factors in the external environment of sperm, this would manifest as an enrichment of adaptively evolving sites in the extracellular regions of the CatSper proteins. We observed different patterns of extra-and intra-cellular changes between the core and auxiliary proteins. Consistent with the patterns observed with CATSPERβ, all of the sites under selection in the auxiliary proteins CATSPERΔ, CATSPERγ, and CATSPERε are in extracellular domains. In contrast, only one third of the sites under selection in the core proteins are extracellular (Supplementary Table 2). These results suggest that in the auxiliary CatSper proteins, adaptive evolution is driven by extracellular interactions. The CatSper complex, therefore, appears locked in an evolutionary conflict that drives its rapid evolution, with *CATSPERβ* being most directly placed at the interface of this conflict.

### The sperm hyper-activation channel Dpkd2 evolves adaptively in Drosophila

Although all *CatSper* genes are missing in *Drosophila*, sperm hyper-activation remains an essential step in successful fertilization (Köttgen et al. 2011). For successful fertilization, *Drosophila* sperm have to swim from the uterus to the sperm storage organs in the female (Neubaum and Wolfner 1999). Several lines of evidence have shown strong selection on different aspects of *Drosophila* reproductive biology, including the length of sperm and ducts (Lüpold et al. 2016), and the peptides that control sperm storage and release (Findlay et al. 2014). In *D. melanogaster*, the gene *pkd2 (Dpkd2*) is required to hyper-activate sperm so that they can navigate the ducts that lead to the sperm storage organs (Gao et al. 2003; Watnick et al. 2003; Köttgen et al. 2011). Like *CatSper, Dpkd2* is a Ca^2+^ channel with a PKD domain. If the CatSper-independent sperm hyper-activation machinery in *Drosophila* is also involved in sexual conflict, we predict to find similar signatures of positive selection in *Dpkd2*.

First, we investigated whether *Dpkd2* evolves under positive selection between *D. melanogaster* and its sister species *D. simulans*. Indeed, *Dpkd2* is notable for having the most significant *p*-value of all genes in the *Drosophila* genome in previous McDonald-Kreitman comparisons between *D. melanogaster* and *D. simulans* (Begun et al. 2007). To conduct more detailed analyses, we performed direct Sanger sequencing of *Dpkd2* from eight strains of each species (Supplementary Table 3). While *Dpkd2* is highly polymorphic within populations, there is also a significant excess of fixed non-synonymous differences between species (McDonald and Kreitman 1991)(Figures 2A and 2B). The fixed non-synonymous changes form discrete clusters. Both the polymorphic and fixed non-synonymous differences are located mostly outside of the PKD domain, indicating the channel pore function may be well conserved while channel activity modulating sites in *Dpkd2* change rapidly.

**FIG 2.**
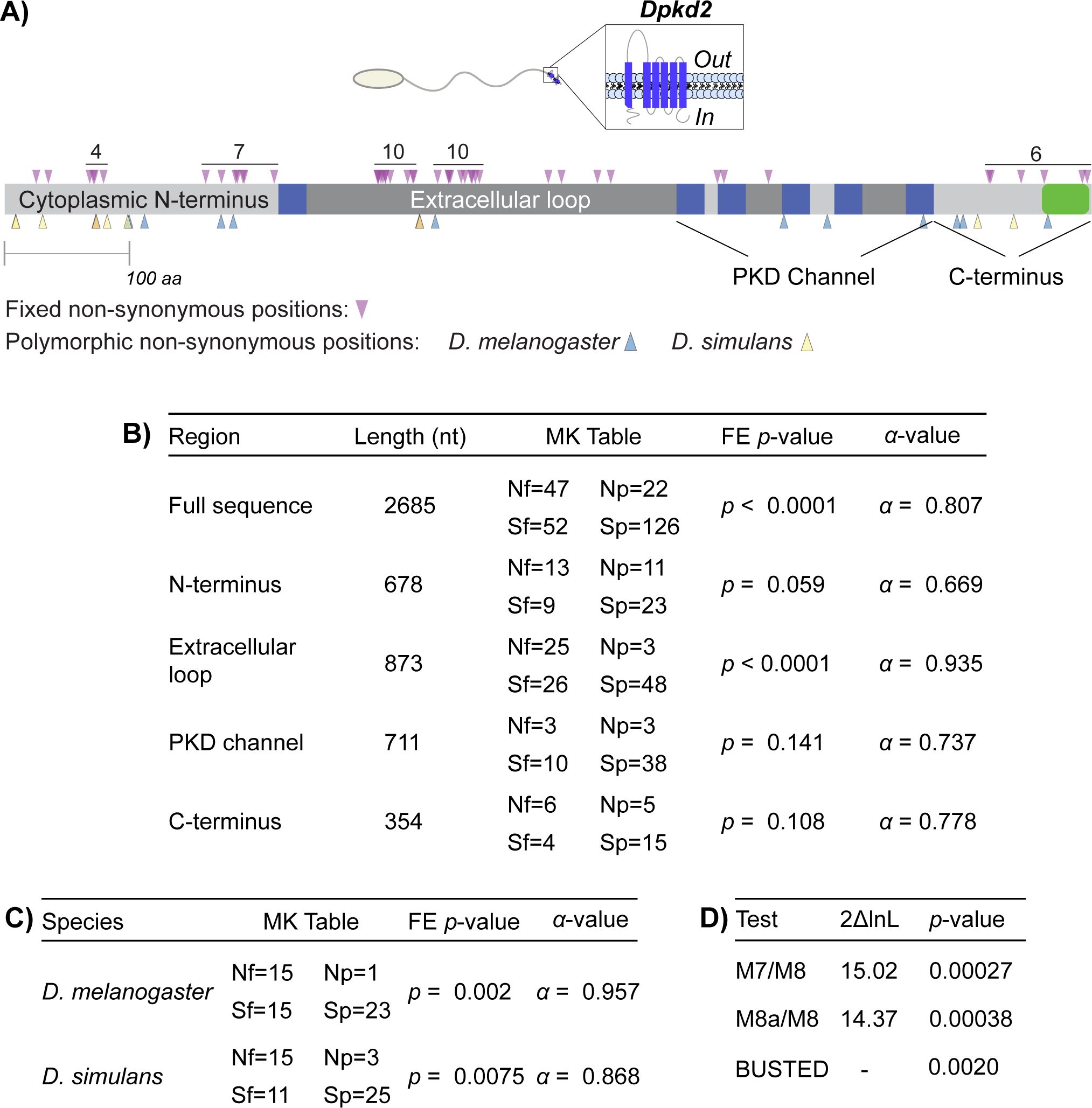
*pkd2* evolves adaptively in *Drosophila*. A) Fixed non-synonymous fixed differences between *D. melanogaster* and *D. simulans* are marked with purple arrows above the domain structure. Clusters of fixed non-synonymous changes are labeled with a bar specifying the number of sites. Polymorphic non-synonymous changes within each species are marked below the domain structure. The gene span of *Dpkd2* is annotated as a grey bar with blue rectangles marking the transmembrane domains and a green box marking the coiled-coil domain. Extracellular and intracellular domains are in different shades of grey. B) McDonald-Kreitman tests show that *Dpkd2* evolves under positive selection between *D. melanogaster and D. simulans*, and this signal is generated by changes in the extracellular domain. The MK table details non-synonymous fixed (Nf), synonymous fixed (Sf), non-synonymous polymorphic (Np), and synonymous polymorphic (Sp) sites. We report the Fisher’s exact (FE) *p*-value, and the *alpha*-value for each segment of the gene. C) A polarized McDonald-Kreitman test for the extracellular domain of *Dpkd2* demonstrates that this region evolves under positive selection along both lineages. Fixed changes were polarized to the *D. melanogaster* or *D. simulans* lineages using *D. yakuba* as an outgroup species. D) NSsites model tests using PAML show that *Dpkd2* evolves adaptively across many species of *Drosophila*.

Many of the fixed differences in *Dpkd2* are clustered in the first extra-cellular loop. A recently solved cryo-electron microscopy (cryo-EM) structure of human PKD2 found that this extracellular loop is critical for gating the homo-tetrameric PKD2 channel (Shen et al. 2016). The orientation of the channel exposes this loop to the external environment of the sperm. To see if *Dpkd2* has evolved adaptively in the extracellular loop, we separately analyzed *Dpkd2* in four regions. Analyses of these separate regions show that it is this extracellular loop that drives the signature of positive selection in *Dpkd2*. To further identify the lineages that experienced adaptive evolution, we polarized our MK test using *D. yakuba* as an outgroup species. Our results show that *Dpkd2* underwent adaptive evolution along each of the lineages that lead to *D. melanogaster* and *D. simulans* (Figure 2C).

To better understand the special positioning of the fixed non-synonymous sites, we modeled a three-dimensional molecular structure of *Drosophila pkd2* with the i-Tasser software (Roy et al. 2010) (Figure 3A). We used the cryo-EM structure of human PKD2 (Shen et al. 2016) as a template, which includes the region from the first transmembrane domain to the end of the Ca^2+^ channel domain. A plot of the non-synonymous changes between *D. melanogaster* and *D. simulans* on this predicted structure shows that these sites are not buried in the pore, but are instead directly accessible to the environment of the sperm (Figure 3B).

**FIG 3.**
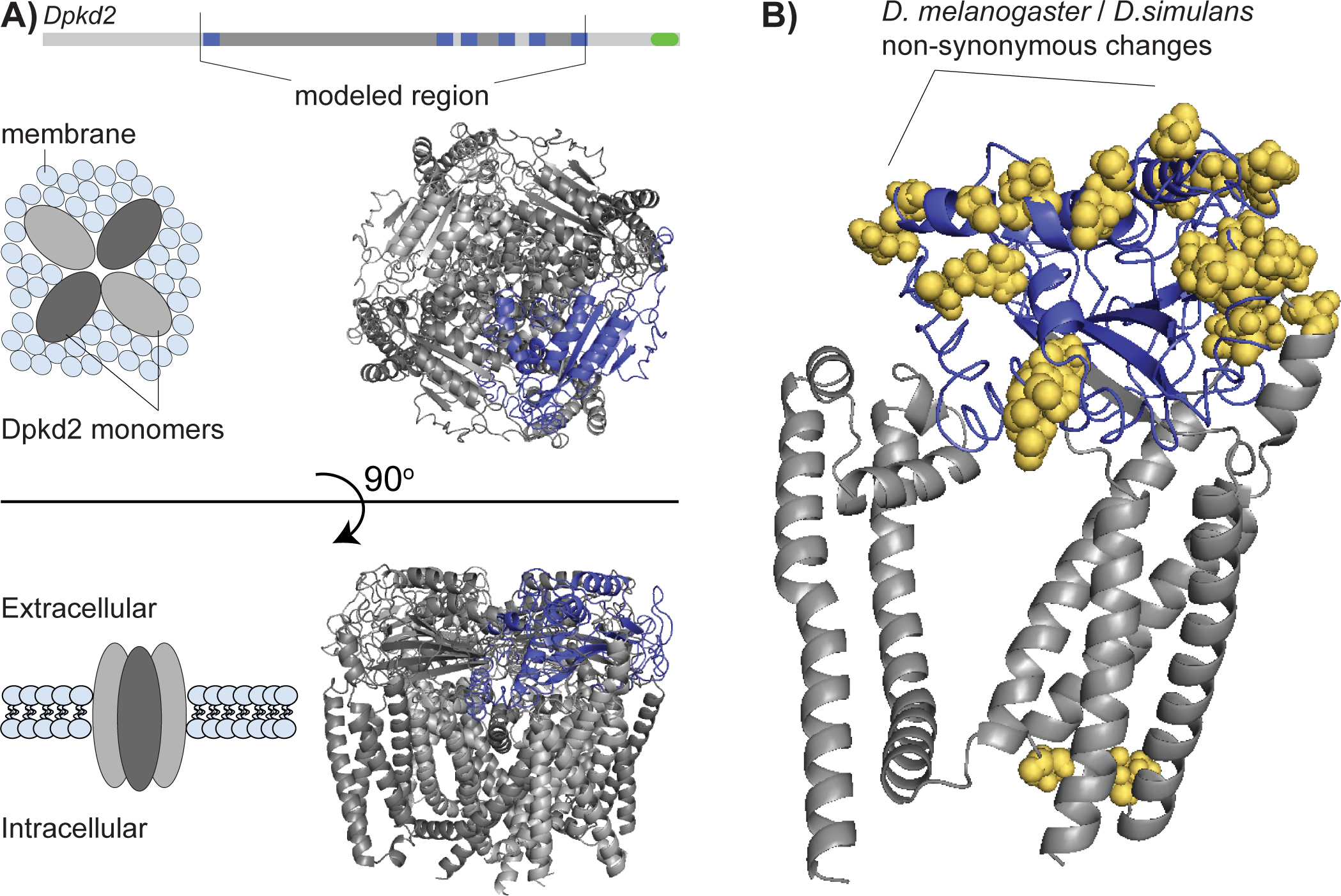
A predicted structural model of *Drosophila pkd2* shows that non-synonymous changes between *D. melanogaster* and *D. simulans* reside on the extracellular faces. A) The homo-tetramer of our predicted *Drosophila* pkd2 structure. The 2-D gene diagram is shaded for the region that could be successfully modeled. The monomers are alternated in shades of grey for contrast. The extracellular loop of one monomer is colored blue. The diagram to the right describes the orientation of the channel. B) Sites that diverge between *D. melanogaster* and *D. simulans* are on the extracellular region of the channel. The base of the monomer is grey, and the extracellular region is blue. The non-synonymous changes between the two species shown in yellow.

Because our results provide strong evidence that *Dpkd2* evolves under positive selection between *D. melanogaster* and *D. simulans*, we next investigated whether a similar signature of recurrent positive selection is seen across a broader range of *Drosophila* species. We curated sequences for *Dpkd2* from 17 species of *Drosophila* by identifying homologs from reference genomes and by directly Sanger sequencing *Dpkd2* from additional species. We used the branch and NSsites models of PAML, and BUSTED to test for signatures of selection in *Dpkd2* (Figure 2D, Supplementary Figure 3) (Murrell et al. 2015). Our results from NSsites and BUSTED analyses provide evidence for episodic positive selection on *Dpkd2* across *Drosophila* species.

Indel variation is known to drive the rapid diversification of the first exon of *CATSPER1* (Podlaha and Zhang 2003). Because there are several fixed indels between *D. melanogaster* and *D. simulans*, we wanted to know if the variation in the length of the extracellular region is greater than that in the other domains of *Dpkd2*. We compared the variance in length of each segment of *Dpkd2* to the variance in length of the whole gene. We find that most of the change in the length of *Dpkd2* comes from the extracellular domain (Supplementary Figure 4). Together, these results show that, similar to *CatSper* genes, both amino acid substitutions and indel differences in regions exposed to the extracellular environment have played an important role in the rapid evolution of *Dpkd2* under recurrent positive selection.

### The sperm function of pkd2 drives its positive selection

In primates, *PKD2* is primarily known for its role in autosomal dominant polycystic kidney disease (Cai et al. 1999). Loss of function in primate *PKD2* causes male sterility, but this sterility is due to abnormal formation of cysts within the testes rather than a sperm hyper-activation defect as in *Drosophila* (Nie and Arend 2014). Because *PKD2* is required for both somatic and gamete development, it is unclear if its evolution would also be driven by sexual selection. Therefore, we analyzed primate PKD2 to see if we could find patterns of selection similar to *Drosophila pkd2* and *CatSpers*.

We first confirmed that primate PKD2 and *Drosophila pkd2* are true orthologs by constructing a maximum-likelihood phylogeny using sequences from the primate *PKD2* family, primate *PKD1* family, *CatSpers*, and *Drosophila pkd2* (Supplementary Figure 5A). Next, we examined the molecular evolution of primate *PKD2* by collecting sequences from 11 primate species and using the same PAML based approach as with the *CatSper* complex and *Drosophila pkd2*. Consistent with our predictions, neither the branch analysis nor the NSsites models suggest any pattern of positive selection (Supplementary Table 5, Supplementary Figures 5B and 5C). Like CatSper proteins, primate PKD2 is part of a complex that contains several other PKD proteins (Tsiokas et al. 1997). We collected sequences and analyzed PKD1, PKD1L1, PKD1L3, PKD2, PKD2L1, and PKD2L2 by the same tests (we did not obtain enough homologous sequences for *PKD1L2* to complete our analysis). We found no evidence for positive selection in primate PKD genes, with the exception of *PKD1L3* (Supplementary Table 5). Interestingly, while the rest of the PKD genes are expressed primarily in the heart and kidney, transcript profiling indicates that *PKD1L3* is most strongly expressed in the placenta, suggesting that it may be subject to a different set of selective pressures than the other PKD genes (Li et al. 2003). Nevertheless, our data show that *PKD2* is well conserved in primates, and the adaptive evolution of *pkd2* in *Drosophila* is likely driven by its role in sperm hyper-activation.

## Discussion

Hyper-activation of sperm, triggered by the opening of flagellar Ca^2+^ channels, is a critical behavioral switch in the race to fertilization. Our analyses show that these Ca^2+^ channels are not conserved, but rather are shaped by recurrent bouts of positive selection. These findings have several implications for understanding the evolutionary and mechanistic aspects of sperm behavior.

First, while much attention has focused on the evolution of the first exon of *CatSper1*, this region reflects only a small part of a much larger pattern. Our analyses show that the entire CatSper complex, including all core and auxiliary proteins, show robust signatures of positive selection in primate lineages. By viewing the molecular evolutionary patterns of the *CatSper* complex as a whole, we find that *CATSPERβ* is the most prevalent target of selection. Little is know about the molecular function of CATSPERβ, and our results suggest that it plays a key role in the regulation of channel activity. Because we find the majority of positively selected sites in the extracellular region of CATSPERβ and the other auxiliary proteins, it is likely that interactions with proteins in the seminal fluid or the female reproductive tract modulate CatSper activity.

Second, we find that the non-orthologous sperm hyper-activation Ca^2+^ channels in primates and flies, taxa that are separated by more than 500 million years, experience remarkably similar selective pressures. We find that the positively selected sites in *Drosophila pkd2* are not buried within the pore, but instead interface with the external environment. Similar to the *CatSper* complex, our results predict that the forces that drive changes in the *Dpkd2* complex involve interacting proteins that modulate the activity of the sperm hyper-activation machinery. These parallel patterns suggest that the sperm hyper-activation machinery may be engaged in the same evolutionary conflicts in a broad diversity of taxa.

Both sperm competition and female choice have the potential to engage the sperm hyper-activation machinery in sexual conflict. Males may deploy seminal fluid peptides to inactivate the sperm hyper-activation channels of competing sperm, thus providing a massive advantage for their own sperm (Figure 4A) (Neubaum and Wolfner 1999; Qazi and Wolfner 2003). Alternatively, females may secrete peptides to inhibit sperm hyper-activation, thus providing a mechanism to modulate fertilization rates and avoid polyspermy (Figure 4B) (Aagaard et al. 2013). Under either scenario, the evolutionary arms race between secreted reproductive proteins and sperm hyper-activation channels drives the patterns of recurrent positive selection that we observe in *CatSpers* and *Dpkd2*.

**FIG 4.**
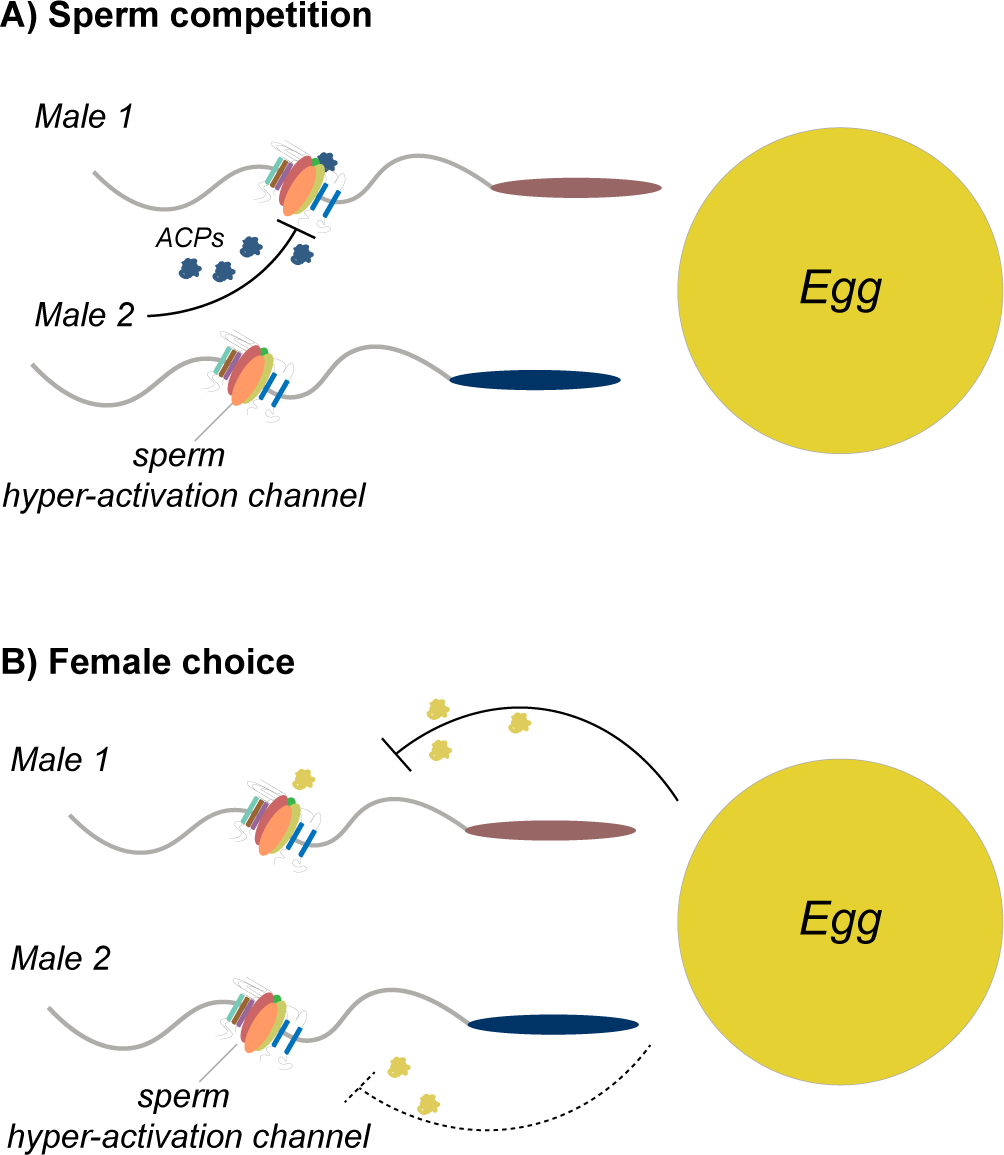
Both inter-male sperm competition and female choice can drive the rapid evolution of the sperm hyper-activation channels. A) Males that secrete seminal peptides that block the hyper-activation channels of sperm from competing males gain a selective advantage. B) Females may secrete peptides that inhibit sperm hyper-activation channels to control fertilization rates. Males that can prevent the inhibition of sperm hyper-activation gain a selective advantage.

Third, the molecular mechanisms of sperm competition remain unclear despite ample evidence for genetic variation in sperm competitive abilities both within and between species (Price et al. 1999; Matute and Coyne 2010; Sweigart 2010; Castillo and Moyle 2014). The patterns of recurrent positive selection on *CatSper* and *Dpkd2* complexes make them strong candidates for a role in sperm competition. While testing the role of *CatSpers* in sperm competition in primates has obvious experimental limitations, testing the role of *Dpkd2* in sperm competition in *Drosophila* is imminently feasible. Our results set the stage for experiments involving transgenic allelic swaps between *D. melanogaster Dpkd2* and divergent *Dpkd2* alleles from other species in an otherwise *D. melanogaster* genetic background. If the sperm of individuals bearing these inter-species allele swaps compete poorly with other sperm, this may reveal an important role for *Dpkd2* in the molecular mechanisms of sperm competition.

Together, our study introduces the sperm hyper-activation genes as a new class of male reproductive proteins that evolve rapidly. We find parallel patterns of adaptive evolution in non-orthologous proteins that serve as the sperm hyper-activation calcium channels in primates and flies. Identifying the factors that modulate the sperm hyper-activation machinery promises to provide insights into the molecular mechanisms used by the sexes to manipulate sperm behavior to their own advantage.

## Materials and Methods

### Collection of primate *and Drosophila* gene sequences

We acquired sequences from up to 17 species of primates and flies each for our phylogenetic analyses. Previous methods for finding homologs for phylogenetic analyses have relied on direct sequencing or manual curating of annotated genomes. Because a reliance on gene annotation models is prone to errors due to mis-annotation, and also limits the number of species that may be analyzed (Markova-Raina and Petrov 2011), we used a different approach to scan sequenced genomes and identify high quality sequences. This method is similar to the one used recently to study positive selection in the synaptonemal complex in *Drosophila* (Hemmer and Blumenstiel 2016), and utilizes the broad range of available sequenced genomes to maximize the power of our analyses. First, we gathered the annotated *H. sapiens* CDS sequence for each *CatSper* gene to use as bait for the searches. For each *CatSper* gene, we gathered homologous sequences from up to 17 primate species that provide a well distributed phylogenetic sampling (Table 1A). Next, we identified the genomic region containing each gene using tBlastn (NCBI) (Madden 2003). Using Exonerate (Slater and Birney 2005) on the narrowed genomic region, we identified the homologous coding regions for each gene, and predicted intron-exon splice sites. We used this two-step method because while exonerate is highly accurate at identifying homologs, it is slow at searching through full genomes. Focusing on smaller contigs that contained our gene of interest accelerated our analyses.

For analyses with *Drosophila* species, we restricted our survey of *Dpkd2* sequences to the melanogaster group of species because including more divergent sequences saturated dS and raised the rate of false positives. For species with annotated genomes, we acquired homologous sequences from Flybase, whereas for species with un-annotated genomes, we used our two-step method homology search (Table 1B). We also PCR amplified, sub-cloned and directly Sanger sequenced *Dpkd2* from *D. tessieri, D. santomea, D. mauritiana*, and *D. orena*.

### Tests of recurrent positive selection

We developed a pipeline to obtain high quality sequences from available genomic resources, and to modify our analysis to ignore low quality sequences (Supplementary Figure 6). Codon based methods of phylogenetic analysis require the accurate alignment of homologous gene sequences. We, therefore, only included sequences from a species when the homolog accounted for at least 90% of the total length of the reference coding region. This method ensured that we did not lose power in our tests because of incomplete gene sequences. To detect patterns of recurrent positive selection in these genes, we prepared and analyzed sequences with the following pipeline. First, we translated all coding sequences to amino acid sequences and aligned the amino acid sequences using CLUSTAL Omega (Sievers et al. 2011). Next, we back-translated the amino acid sequences to their corresponding CDS using Pal2Nal (Suyama et al. 2006), using settings to remove all gaps and stop codons. We then manually reviewed each alignment to confirm that there were no gaps or stop codons. We constructed the phylogenies for our alignment based on modEncode data (Perelman et al. 2011; Chen et al. 2014).

We used the same alignments for two separate analyses: Phylogenetic Analysis by Maximum Likelihood (PAML) (Yang 2007) and Baysean Unrestricted Test For Episodic Diversification (BUSTED) (Murrell et al. 2015). To test for recurrent selection using PAML, we compared NSsites models M7 and M8, using the branch model 0 and the standard clock. We calculated a *p*-value using a log-ratio test between the log-likelihood scores for each model (Yang 2007). PAML is highly sensitive to misalignments; even slight mis-alignments can easily create false positive signals (Markova-Raina and Petrov 2011). When we observed a *p*-value < 0.05, we repeated our analyses by starting at the beginning of the pipeline, but this time using the T-coffee (Notredame et al. 2000) and Muscle (Edgar 2004) aligners, ensuring that our result was not an artifact of alignment error. We only considered true rejections of the null where a *p*-value < 0.05 was observed with all three aligners, and reported the least significant *p*-value. We ran M8a using the same alignment that generated the least significant *p*-value. Like PAML, BUSTED compares models of selection for homologous sequences over a phylogenetic distribution. Unlike PAML, BUSTED takes a Bayesian approach to build these models. This framework makes BUSTED an independent test from PAML to analyze the molecular evolution of a gene. We ran BUSTED using the Data Monkey server (http://www.datamonkey.org/). When the program did not correctly compute the base tree, we re-oriented nodes to correctly reconstruct the phylogeny.

To compare *dN/dS* between the *CatSper* genes (Figure 1A), we removed the percent alignment threshold for each species so that we could analyze *dN/dS* over a common phylogeny of 16 species. To calculate *dN/dS* along branch lengths, we used the branch model 1 with model 0 (Yang 2007).

We also used the McDonald-Kreitman (MK) test for positive selection on the narrower timescale of *D. melanogaster* and *D. simulans* divergence. For this test, we sub-cloned and re-sequenced *Dpkd2* from eight lines of *D. melanogaster* and *D. simulans* using the method described above. We ran MK tests for *Dpkd2* using DnaSP (Librado and Rozas 2009). To identify protein domains, we submitted amino acid sequences to the SMART server (Letunic et al. 2015) using all available databases.

### Insertion/Deletion Polymorphism Analysis

To quantitatively assess the extent of indel differences between *Dpkd2* in the *D. melanogaster* group, we developed a null prediction for indel variation per amino-acid site. To do this, we first measured the differences in length for *Dpkd2* for the full gene, and determined the length variance per base pair. We measured the difference in length of each region of the gene, using the N-terminus, first trans-membrane domain, PKD channel, and C-terminus as alignment anchors. We then calculated the standard deviation of each of these regions, and divided by the length of the region to calculate a deviation per amino-acid value. For each region, we present the ratio of one deviation per amino-acid of the gene region divided by the full length of the gene.

### Computational modeling of the *Dpkd2* three dimensional structure

To model the structure of *Dpkd2* we accessed i-Tasser via the web server at http://zhanglab.ccmb.med.umich.edu/I-TASSER/ (Roy et al. 2010). We provided the program with the amino acid sequence of *Dpkd2* from positions 233-810 to correspond with the published high resolution cryo-EM structure for human PKD2 (Shen et al. 2016). We provided the human PKD2 structure as a scaffold for i-Tasser, and used the option to align the two sequences before structural prediction. Due to the limits of i-Tasser, we modeled a single monomer of *Dpkd2*. To arrange these monomers in a tetrameric complex, we aligned the *Dpkd2* monomer to the four positions of the human PKD2 monomers in the solved structure using PyMOL. We highlighted all non-synonymous positions between *D. melanogaster* and *D. simulans* in the predicted structure.

## Acknowledgements

This work was supported by the National Institutes of Health (Developmental Biology Training Grant 5T32 HD0741 [to JCC]; R01 GM115914 [to NP], a Mario Capecchi endowed assistant professorship (NP), and the Pew Biomedical Scholars Program. We thank Nora Brown, Chris Leonard, and Mia Levine for helpful comments with this manuscript.

